# Cell Spotter (CSPOT): A machine-learning approach to automated cell spotting and quantification of highly multiplexed tissue images

**DOI:** 10.1101/2023.11.15.567196

**Authors:** Ajit J. Nirmal, Clarence Yapp, Sandro Santagata, Peter K. Sorger

**Author notes:** Equal contribution (Ajit J. Nirmal, Clarence Yapp).

## Abstract

Highly multiplexed tissue imaging and in situ spatial profiling aim to extract single-cell data from specimens containing closely packed cells of diverse morphology. This is challenging due to the difficulty of accurately assigning boundaries between cells (segmentation) and then generating per-cell staining intensities. Existing methods use gating to convert per-cell intensity data to positive and negative scores; this is a common approach in flow cytometry, but one that is problematic in imaging. In contrast, human experts identify cells in crowded environments using morphological, neighborhood, and intensity information. Here we describe a computational approach (Cell Spotter or CSPOT) that uses supervised machine learning in combination with classical segmentation to perform automated cell type calling. CSPOT is robust to artifacts that commonly afflict tissue imaging and can replace conventional gating. The end-to-end Python implementation of CSPOT can be integrated into cloud-based image processing pipelines to substantially improve the speed, accuracy, and reproducibility of single-cell spatial data.

## MAIN

Recent advances in using highly multiplexed imaging to analyze normal and diseased tissues have made it possible to map cell types and states in preserved 2D and 3D environments^1,2^. These technologies enable the quantification of 20-80 different protein antigens at subcellular resolution using either sequential (cyclic)^3–7^ or one-shot data acquisition^8,9^. A subset of the antigens in these images are used to identify cell type (e.g., CD4 immune cells) whereas others measure features of cell state such as response to cytokines (e.g., IRF transcription factors), cell cycle state (e.g., Ki67 or PNCA), or juxtracrine signaling (e.g., PD1 and PDL1). A whole-slide image of several square centimeters comprises 10^6^ to 10^7^ cells and cannot be fully understood by human inspection alone. Moreover, integration of high-plex imaging with other types of single-cell data, including scRNA-SEQ and spatial transcriptomics, requires processing images to extract quantitative data on individual cells. This typically involves computationally assembling (i.e., stitching) individual image tiles and channels into a composite high-plex image, finding the positions of individual nuclei, segmenting the image to determine the borders of individual cells, quantifying the intensities of protein antigens within each cell, writing these metrics to a “spatial feature table” in which each row represents a single cell, and then processing the resulting table to identify cell types, states, and interactions^10^.

Although significant effort has gone into developing automated methods for single-cell analysis using high-plex images, the task remains inherently challenging. The close packed nature of tissues often involves cells with irregular and intertwined processes, complicating segmentation^11^. Additionally, immunofluorescence-based images typically display a relatively poor signal-to-noise ratio at the level of individual pixel intensities with much of the information lying in morphology, adding to the complexity of analysis. While a flow cytometer gathers intensity data by consolidating the fluorescent signal from a cell into a single data point, tissue imaging microscopes having a resolution of ∼400 nm (e.g., using a 20X high performance objective) distribute intensity across 200-400 pixels. This dispersal leads to a lower signal-to-noise ratio in each pixel but also enables microscopes to excel at recording the structure and spatial organization of cells.

The task of gating high-plex images becomes more challenging as dataset size increases, which is in critical for drawing conclusions with robust spatial power^12,13^. A recent study involving 75 whole-slide images required gating ∼10^8^ cells across 18 fluorescence channels and was both slow and potentially error prone. In principle, subtracting a no-antibody control image from one stained with a fluorescence antibody helps mitigate this problem, as do image processing methods that identify and eliminate artefactual and outlier signals. However, these approaches typically fall short in accurately identifying true signals across an entire image due to the irregularity of the background across the large number of channels in a high-plex image.

We have found that differentiating background noise from true signal requires the greatest level of human intervention. Gating tools derived from flow cytometry, or with integrated image viewers such as Gater (https://github.com/labsyspharm/minerva_analysis), allow users to set gates on one- or two-channel marker distribution plots (a process akin to flow cytometry) while visually reviewing the underlying image. In this case, the human expert uses morphological, and neighborhood information to set an intensity gate, but this is time-consuming and subjective. Semi-automated techniques such as 2-class Gaussian mixture modeling^14^, in which one Gaussian represents the true signal and the other the background, can be used to estimate gate values, but these commonly require human refinement. To eliminate the need for manual assessment, machine-learning (ML) approaches have emerged for direct cell type assignment^15–18^, however, these methods often fail, particularly when images have one or more channels with high background. Moreover, automated cell type assignment does not always apply to markers that measure cell state (as opposed to cell type); this is particularly true of antigens whose levels vary continuously from one cell to the next and cannot easily be discretized.

In this paper, we investigate the factors that make it challenging for both humans and algorithms to gate intensity data in highly multiplexed images and describe the development of ‘Cell Spotter’ (CSPOT) for automated detection of true positive cells. CSPOT is an early example of an image processing tool that operates at both the pixel and single-cell level to extract data from tissues containing many different types of closely packed cells. CSPOT uses a two-stage process that starts by training a deep learning (DL) model at the pixel level from a human-curated (supervised) set of true positive and true negative cells. A series of ML algorithms then leverages the probability scores from the DL model and raw intensity values computed at the level of single cells to accurately score all positive and negative cells in an image. CSPOT thereby mimics the ability of humans to jointly evaluate morphology, intensity, and neighborhood in making call assignments, and we demonstrate that CSPOT is robust to a wide range of artefacts that interfere with conventional intensity-based gating.

## RESULTS

We used high plex images from two previously described datasets: (i) a 64-plex CyCIF image of a tissue microarray (TMA) containing 123 cores from 34 different tissue and tumor types^10^; each core is 0.6 mm in diameter and contains an average of 10,500 segmented cells, (ii) a 24-plex 3D CyCIF dataset of a human colorectal adenocarcinoma (CRC) resection^12^ consisting of 25 five sections each 5 µm thick and∼ 4-5 cm^2^ in area spaced 25 µm apart along the Z axis; the dataset comprises ∼ 5 x 10^7^ cells (see **Supplementary Table 1** for HTAN ID numbers and data access). We used the TMA data to test methods across tissue and cell types in a dataset with a relatively high artifact level; the CRC data provided a setting in which to test performance on a good quality whole-slide image. Both datasets had previously been processed using the MCMICRO pipeline^10^ to generate stitched images that were aligned across channels (i.e., registered), segmented for individual cells, and quantified for single-cell features. The output of MCMICRO is a spatial feature table that includes the mean fluorescence intensity (MFI) for each marker integrated over the segmentation mask and the position in space of each cell in the image.

To better understand the gating process in the context of high-plex images we compared four existing methods: (i) manual gating on an MFI frequency plot; (ii) two-class Gaussian Mixture Modeling (GMM) on the MFI plot (under the assumption that one Gaussian distribution corresponds to background and the second to positive signal); (iii) manual bi-marker gating as commonly performed for flow cytometry data and; (iv) interactive image-based gating in which a human annotator adjusts the gating threshold while visually inspecting the underlying images for morphological, neighborhood, and intensity information. Although laborious, this approach was judged the most accurate based on an informal assessment of inter-observer agreement and was used to generate ground truth annotations for evaluating the other methods.

Staining of cell membranes by E-cadherin (ECAD; a marker of epithelial cells) in colon tissue is an example of straightforward manual gating: the MFI frequency plot is clearly bimodal, with the lower intensity mode corresponding to diffuse background signal and the high intensity mode corresponding to epithelial cells exhibiting the characteristic morphology of colonic villi. In this case, an unambiguous threshold could be set (dotted line) to distinguish ECAD positive from negative cells (**Fig. 1a,b**). In contrast, the pan-immune cell marker CD45 exhibited a single low intensity mode and an extended rightward shoulder (**Fig. 1c**). Using a bi-marker gating approach with ECAD as the second (orthogonal) marker, it was possible to delineate a threshold (dotted line) that demarcated the CD45-positive cells (**Fig. 1d**). Subsequent visual review showed that cells residing in the extended shoulder had the characteristics of immune cells (**Fig. 1e**). Note, however, that no cell should stain positive for both ECAD and CD45 and datapoints having this property represent errors in segmentation^11^. Compared to manual gating, using a two-class Gaussian Mixture Model (GMM) omitted up to 40% of all true positive CD45^+^ cells, as judged visually and by using bi-marker gating (**Fig. 1f**). In our experience, substantially more antibodies in high-plex tissue images are associated with unimodal intensity profiles with shoulders (like CD45) as opposed to bimodal profiles (like ECAD), demonstrating the general challenge associated with setting intensity gates. We therefore sought to develop an ML framework that, like human annotators, uses morphological information to identify positive and negative staining cells.

**Fig. 1:**
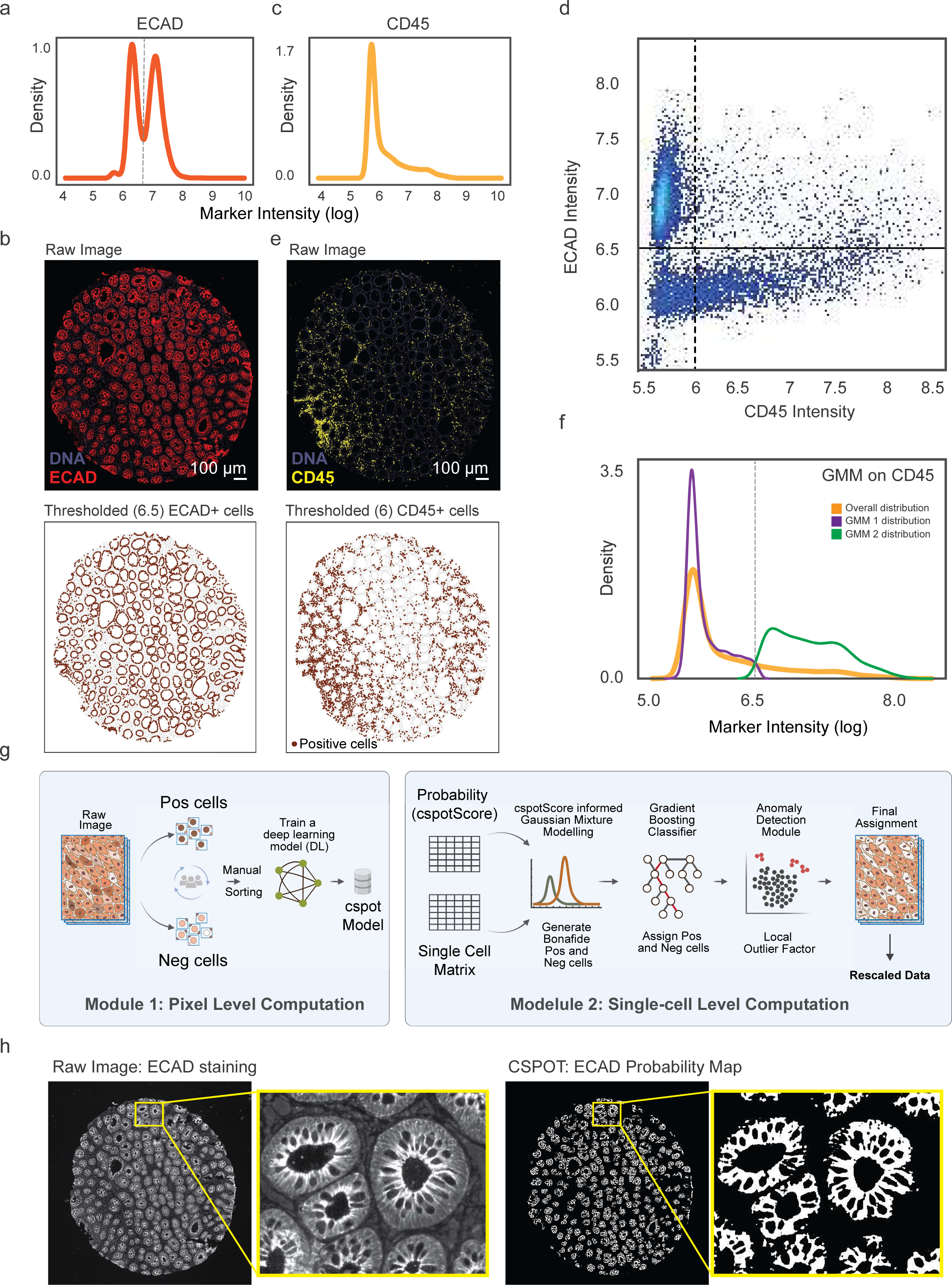
Common approach to differentiating signal from background. a, Distribution plots of ECAD intensity from a colon sample core, with x-axis representing marker intensity values in log scale and y-axis representing the frequency of occurrence. b, Top: Image of ECAD staining of a colon sample. Bottom: Dot plot representing cells with an ECAD intensity greater than 6.5 in brown. c, Distribution plots of CD45 intensity from a colon sample core, with x-axis representing marker intensity values in log scale and y-axis representing the frequency of occurrence. d, Two-dimensional density plot, where the intensity of the CD45 marker is represented on the x-axis, and the intensity of ECAD is represented on the y-axis. e, Top: Image of CD45 staining of a colon sample. Bottom: Dot plot representing cells with a CD45 intensity greater than 6 in brown. f, Distribution plots illustrating the intensity of CD45 (orange) in a colon sample core. The violet and green distributions correspond to the two Gaussian distributions identified by applying a 2-class Gaussian Mixture Modeling prediction to the data. g, Schematics of the CSPOT algorithm depicting the steps involved in pixel and single-cell level prediction. h, Left: The overall and zoomed-in views of ECAD staining in a colon sample. Right: The CSPOT prediction of ECAD in the same sample.

### Overview of CSPOT Algorithm

CSPOT consists of two modules (**Fig. 1g**). Module 1 employs a U-Net Convolutional Neural Networks (CNN), trained on user-curated true positive and negative examples, to perform pixel-level semantic segmentation, assigning each pixel a probability score per marker. Module 2, at the single-cell level, utilizes the probability scores along with single-cell MFI data (spatial feature table) to identify true positive and negative cells. CSPOT includes a training phase and a prediction phase; Module 1 is used in both, while Module 2 is used only in prediction.

During training, Module 1 generates 64x64 pixel thumbnails centered on cell nuclei. These thumbnails are sorted by experts into high confidence sets of either true positive or true negative staining (**Extended Data 1a**); ambiguous cases are omitted. Next, training labels are automatically generated using the OTSU thresholding technique, creating binary masks for each thumbnail. These masks, along with standard image augmentations such as rotation, flipping, dropout, and L2 regularization, are employed to train a 2-class, 4-layer U-Net CNN with 16 input filters. To further enhance the model’s ability to learn cellular morphology, we implemented intensity scaling augmentation (detailed in **Methods**).

During prediction, unseen images are input into Module 1, generating pixel-level probability masks with scores between 0-1, representing the likelihood that a pixel represents a true positive for a given fluorescent marker (**Fig. 1h**). These scores are aggregated to yield a median *cspotScore* for each cell. In the next step, Module 2 uses a two-class GMM to sort cells into either positive or negative categories based on the *cspotScore*. The mean MFI values from this sorting are then used to set up another similar analysis, but this time on the raw MFI data (spatial feature table). This second analysis provides another set of categorized cells as either positive or negative. A common set of positive and negative cells is then identified from the intersection of both. This common set of *bona fide* positive and negative cells is used to train a Gradient Boosting Classifier^19^ to predict marker positivity across the entire image. Predictions are refined using a Local Outlier Factor (LOF) algorithm^20^ for anomaly detection (e.g., a PanCK^+^ epithelial cell that clusters away from other predicted PanCK^+^ cells). CSPOT’s final predictions convert marker intensity to a normalized score via a sigmoid function, scaling intensities to between 0-1. The final scaled values preserve the independent distributions of negative and positive cells in the original data and values above 0.5 are considered positive cells.

CSPOT is available as a standalone Python package compatible with a command line interface and interactive modes (e.g., with Jupyter notebooks). It’s Dockerized for easy integration into cloud-based pipelines such as MCMICRO^10^. The processing time for generating probability masks (module 1) varies by image size and GPU availability, but the remainder of the algorithm (module 2) takes approximately one minute per marker to process 10^6^ cells on a standard laptop (**Extended Data 1b**). When applied to ECAD and CD45 CyCIF data from **Fig. 1**, we observed effective discrimination of cells positive for either ECAD or CD45 in line with manual gating (**Extended Data 1c**). We also confirmed cross-platform compatibility using two commercial multiplexed imaging platforms: COMET (from Lunaphore Technologies SA), which uses sequential immunofluorescence with primary and secondary antibodies, and CODEX (e.g., on instruments from Akoya Inc.), which uses DNA-barcoded antibodies. Models trained on CyCIF images successfully predicted CD45^+^ and CD3D^+^ cells in COMET data (**Extended Data 1d)** and CD8^+^ cells in CODEX data (**Extended Data 1e**).

### Performance of CSPOT at the single-cell level

To evaluate the performance of CSPOT, we compared it to expert human annotations made on four TMA cores comprising colon, seminoma (a germ cell tumor) and two lung mesothelioma samples (4.9 x 10^4^ cells total) across six markers (ECAD, CD45, CD4, CD3D, CD8A, and KI67; **Fig. 2a**). The model training for these markers was performed using an independent colon and tonsil sample. The cores exhibited nuclear staining for the KI67 proliferation marker, cytoplasmic staining for ECAD, and cell surface staining for the immune lineage markers CD45, CD4, CD3D and CD8A in a variety of patterns: (i) strong staining of immune and epithelial cells having the expected morphologies (ii) artefactual staining patterns with irregular background and “streaking” (iii) antibodies that strongly stained one core but not another, likely reflecting absence of the antigen (iv) intermixing of frequently and infrequently positive cells (e.g. for CD45 and Ki67 respectively) and (v) cells with differing degrees of packing and morphologies (e.g. immune cells and ECAD positive epithelial cells in colon tissue). For each of the six markers, manual bi-marker and visual gating were performed by human experts and used as a benchmark for CSPOT performance based on accuracy, precision, and F1. Accuracy is the measure of how closely the predicted values align with the actual or true values. Precision is a metric that quantifies the fraction of true positives within the set of all positive predictions; the greater the precision, the fewer false positives are present within the CSPOT results. The F1 score is a weighted harmonic mean of precision and recall. Excluding KI67 predictions from all cores (see below) CSPOT achieved an average accuracy of 95% across samples and markers with a precision averaging 0.88, F1 scores between 0.85 and 0.98 (depending on sample; **Extended Data 2a**). CSPOT could enumerate relatively rare cell types (2.6% KI67^+^ cells in core 2) as well as abundant close-packed cells (70% CD45^+^ cells in core 4), demonstrating the versatility of the algorithm.

**Fig. 2:**
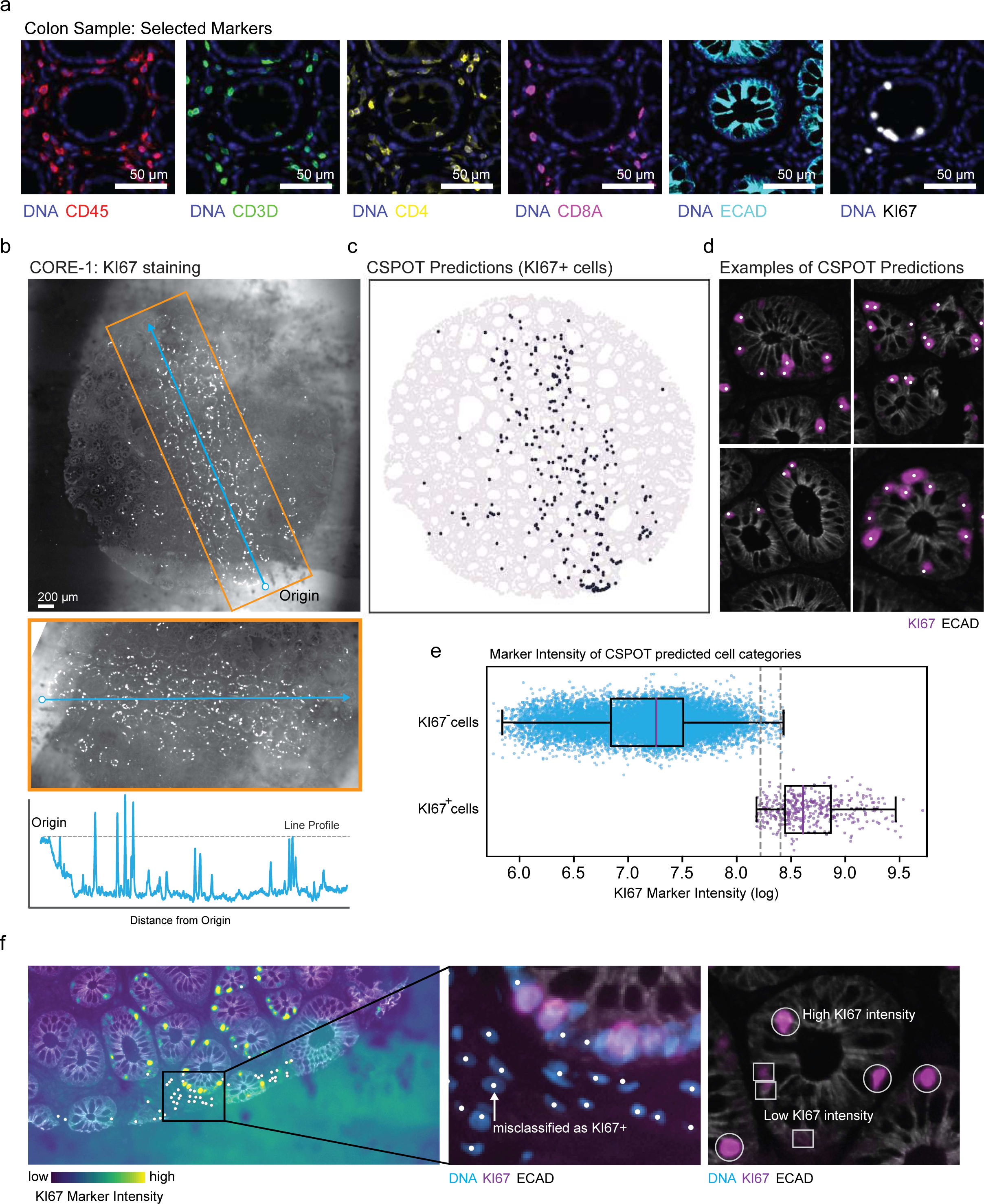
CSPOT accurately assigns marker positivity at the single-cell level. a, Exemplary staining patterns of markers that were selected for in-depth investigation (CD45, CD3D, CD4, CD8A, ECAD, KI67) in colon tissue. b, KI67 staining in a colon sample. Middle: A closer view of the orange box in the top panel. Bottom: A line plot of KI67 marker intensity along the blue line indicated in the above two figures. c, Cells identified as KI67 positive by CSPOT mapped onto the spatial coordinates of the image shown in (b). d, Exemplary regions of the image presented in (b) superimposed with points that were identified as KI67^+^ cells by CSPOT. KI67 staining is represented using the pseudo color violet. e, Box plot of KI67 marker intensity in cells classified as positive or negative by CSPOT. The y-axis represents the two groups of cells, and the x-axis represents the marker intensity values in log scale. Gray dotted lines indicate the range of expression intensities where there is an overlap between negative and positive cells, which could lead to misclassification if a single threshold was used to identify positive and negative cells. f, A tissue region containing cells that fall between the grey dotted lines in figure (e). Left plot shows pseudo-colored KI67 staining of a region where the cells have KI67 intensity higher than the lowest true KI67^+^ cell. Middle plot shows a zoomed-in view of the region boxed in the left panel. Right plot highlights true-positive KI67 cells in a different region that exhibit lower intensities of KI67 than the background noise highlighted in the left panel.

KI67 staining in Core 1 represented a situation where a single gate was ineffective (**Extended Data 2b,c**). When we examined KI67 fluorescence intensity along an arbitrary line through an image of a core (**Fig. 2b**) we found that it varied 2-3-fold at peaks corresponding to foreground intensity (true cells) and that background intensity was highly variable; in some cases, the background was higher than the foreground signal level for true positive cells elsewhere in the image. CSPOT was nonetheless able to correctly identify cells that were KI67^+^ (∼3% of the all cells) including those that resided in a band at the bottom of the intensity scale (in **Fig. 2b**) that comprised a mixture of positive and negative cells (**Fig. 2c,d**). This low intensity KI67^+^ cells corresponded to ∼33% of all KI67^+^ cells in the image, which means that simple gating approaches result in a high level of misclassification (**Fig. 2e**). Inspection of the underlying images confirmed our understanding of why this arises, namely that cells stained either weakly or strongly with Ki67 (a known feature of Ki67 staining^21^) were overlaid on a variable intensity background (**Fig. 2f**). Based on these data, we conclude that CSPOT is superior to GMM and manual gating and similar to interactive image-based gating by humans at identifying cells that stain positive or negative for antigens in the cytoplasm, nucleus, or cell surface in the presence of high and variable background.

### Sensitivity testing

The performance of ML algorithms on images is influenced by the quality and depth of training data and, for complex scenes, by the abundance of specific elements in each scene. To evaluate the sensitivity of CSPOT to the amount of training data, we randomly sampled 3 to 1000 cells positive for a particular marker (e.g., ECAD) from each core and used that to make the final predictions. Random sampling was repeated 100 times, resulting in synthetic cores containing a pre-specified number of positive cells alongside cells positive for all other markers in a proportion typical for that core. We found that a minimum of ∼500 cells were necessary for CSPOT achieve an F1 score of 0.90 across the four cores (**Extended Data 2d**). Since a typical TMA core contains ∼1 x 10^4^ cells and a whole slide image contains ∼2 x10^6^ cells, we estimate that CSPOT can detect cells present at a frequency of 5% in a TMA core and <0.05% in a whole slide image. We also investigated the number of unique images necessary for training a CSPOT model, as assessed by an intersection-over-union (IoU) metric; in this context, IoU measures the overlap in the area between an input mask (auto generated by OTSU) and probability maps generated by CSPOT (thresholded at 0.5). Since the CNN has two classes, we evaluated IoU for both foreground (marker intensity) and background classes. We found that IoU increased until ∼ 200 positive and negative training examples were included in the model (**Extended Data 2e**). In practice, this occupies a single human annotator about one hour per marker to accomplish, which compares favorably to manual gating.

### Overcoming complex background intensities with CSPOT for analysis of multiple tissues

A common application of tissue microarrays (TMA) is to compare cell types or biological processes across a range of normal or diseased tissues. As is common for a TMA with such a wide variety of specimens, some combinations of antibodies and TMA cores in our images exhibited staining patterns that do not resemble cells or known tissue structures. Even when staining appeared to be specific, it varied substantially in intensity from one specimen to the next, likely for a variety of reasons including tissue type-dependent non-specific antibody binding^22^, incomplete washing or partial dissociation of unstable antibody-antigen complexes^23^, differences in tissue fixation time or storage conditions of the tissue blocks, and true biological variation. To evaluate the robustness of CSPOT to these types of problems we evaluated performance on cores selected for evidence of data that was difficult to process using conventional methods.

The first challenge involved identifying true positive cells in a series of cores on the same TMA but for which staining intensities were highly variable. **Fig. 3a** shows colon lymph node and meningioma cores with a 5.2-fold difference in mean anti-CD4 staining. A single TMA-wide intensity threshold failed to discern positive and negative cells across the cores, but a CSPOT model effectively identified CD4^+^ cells in both cores without needing core-specific training data (**Fig. 3b**).

**Fig. 3:**
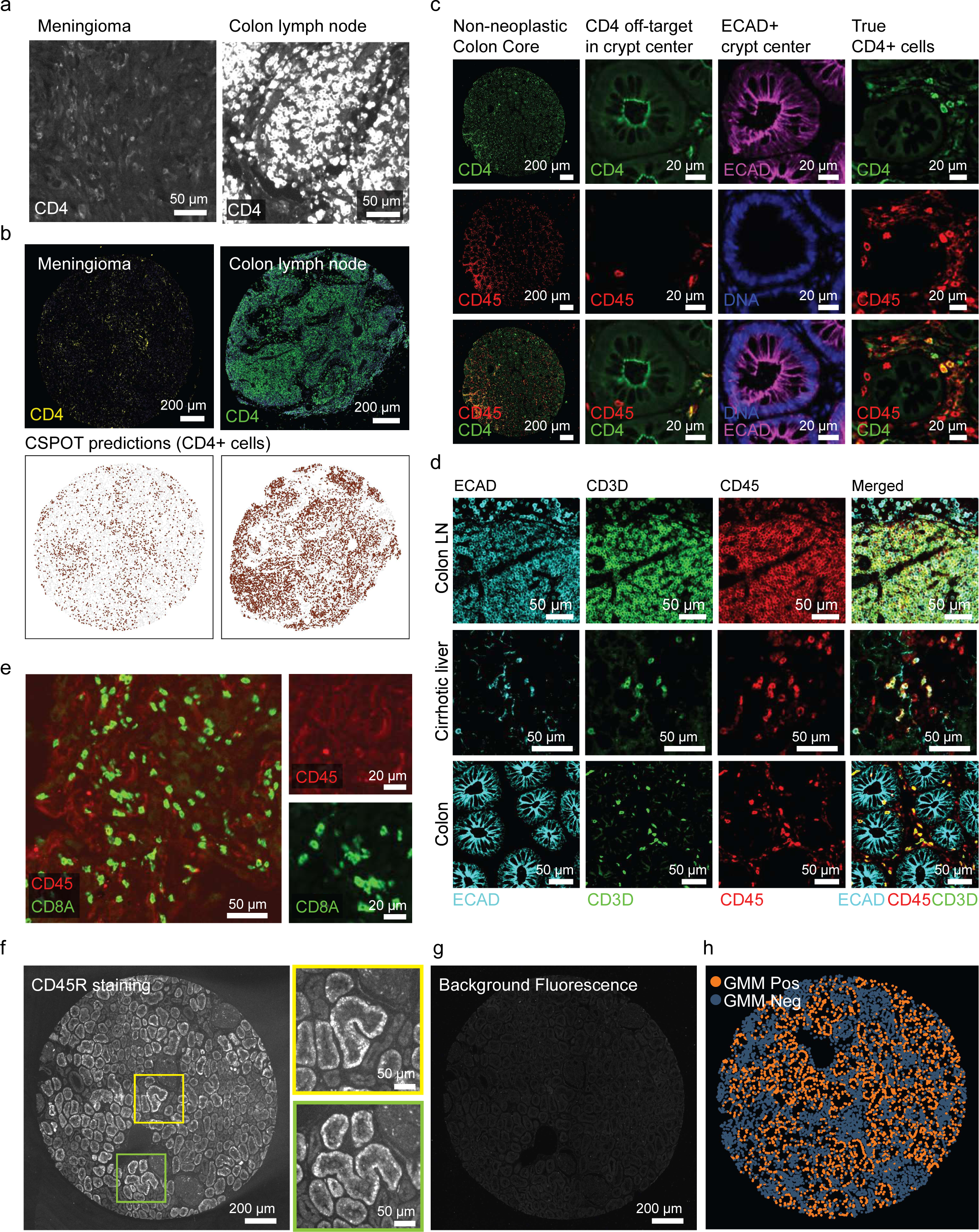
Artifactual staining patterns that impede automated processing of multiplexed imaging. a, Differential staining intensity of CD4 between two cores stained and rendered using the same conditions. b, Top: Zoomed-out view of the samples presented in (a), highlighting contrasting CD4^+^ cell staining. Bottom: The CSPOT predictions of the CD4^+^ cells in the same samples. c, The staining of CD4 (green), CD45 (red), and ECAD (violet) in multiple zoomed-in regions of a colon sample, highlighting the non-specific binding of CD4 on ECAD^+^ epithelial cells towards the center of the colon crypt, and the adjacent regions that exhibit specific staining of CD4 on CD45^+^ immune cells. d, Staining of ECAD (cyan), CD3D (green), and CD45 (red) in three sample (colon lymph node, cirrhotic liver and colon). Top and middle rows highlight the non-specific binding of ECAD on immune cells (CD45/CD3D). Bottom row demonstrates the specific binding of ECAD to epithelial cells, and its absence on CD45/CD3D positive immune cells. e, Left: Image highlighting the loss of CD45 intensity (red) in non-neoplastic colon samples. Right: Magnified views highlighting that some CD8^+^ immune cells (green) do not express CD45 (red). f, CD45R staining on normal kidney cortex. Left: Insets display magnified views of specific regions of the sample that highlight the non-specific binding of antibodies, where staining is not restricted to the cell surface. g, The autofluorescence channel rendered with the same settings as (f), indicating that the high background is not a result of tissue autofluorescence. h, Positive and negative cells predicted by the Gaussian Mixture Modeling (GMM) algorithm for the image shown in (f) and mapped to the spatial coordinates of the image.

A second challenge involved off-target antibody binding in a subset of TMA cores; this is an issue when TMAs arise for a wide range of tissue types. For instance, anti-CD4 antibody exhibited non-specific staining of ECAD^+^ CD45^-^ cells in centers of colonic crypts, despite correctly staining CD45^+^ CD4^+^ ECAD^-^ immune cells nearby (**Fig. 3c**). Similar issues arose with non-specific anti-ECAD binding to CD45^+^ cells in a colon lymph node (**Fig. 3d**) and CD45^+^ CD3^+^ cells in cirrhotic liver (**Fig. 3d**). Additionally, a non-neoplastic lung sample exhibited an unexpected pattern of CD8^+^ T cells lacking CD45 expression (**Fig. 3e**). These issues weren’t universal across the TMA, as most cores showed high antibody specificity. However, problematic cases stood out in computational analysis as outliers in shape and intensity. CSPOT fully resolved the two situations described above, rejecting non-specific CD4 staining in the colonic crypts and ECAD staining in cirrhotic liver. In the case of anti-ECAD antibodies binding to CD45^+^ cells, CSPOT exhibited an 8% error rate (**Extended Data 3a**). Notably, errors were mainly localized along tissue edges, an area requiring further investigation because it has characteristics distinct from the bulk of the specimen. Additional issues were also noted involving the non-specific binding of antibodies, which exhibited variable intensities both across different tissues (**Fig. 3h**; see Supplementary Note) and within individual specimens (**Extended Data 3b-g**; see Supplementary Note).

These examples highlight the extent to which problems with staining can be both antibody and tissue type specific; when a type of cell is found in many tissues and expected to stain in a similar manner (e.g., a T cell) it is easy to confuse positive and negative signals with differences in cell abundance. To study this problem more systematically, we conducted a manual review of five T cell markers (CD45, CD45R, CD3D, CD4, and CD8A) in 123 TMA cores (615 images in all) and identified 158 in which no cells exhibited positive staining for one of the markers but instead involved diverse staining patterns of non-specific and/or high background staining. In comparison to the manual benchmark (at the whole core level), CSPOT achieved an average accuracy of 93%, ranging between 85% to 97% (**Fig. 4a**). To study assignments that differed between CSPOT and human annotators, we constructed a series of confusion matrixes (one per marker; **Fig. 4b**). We found that CD3D exhibited the greatest disparity between CSPOT and human curators (16 mismatched assignments out of 123); when we examined CD3D^+^ cells detected by CSPOT we found that many had low probability scores, leading to low confidence predictions (**Extended Data 4a**). To determine why, we examined CD3^+^ cells in the training data from tonsil and found that cells were clustered closely together (**Fig. 4c**) whereas they were interspersed with other cells in most tissues - a difference that confounded CSPOT but not humans (**Fig. 4d**). When we retrained the CSPOT model for CD3D using examples of T cells that were sparsely distributed (e.g., in colon tissue sample; **Fig. 4e**) accuracy across the TMA increased from 85% to 96%, which raised overall accuracy across all markers to 96%.

**Fig. 4:**
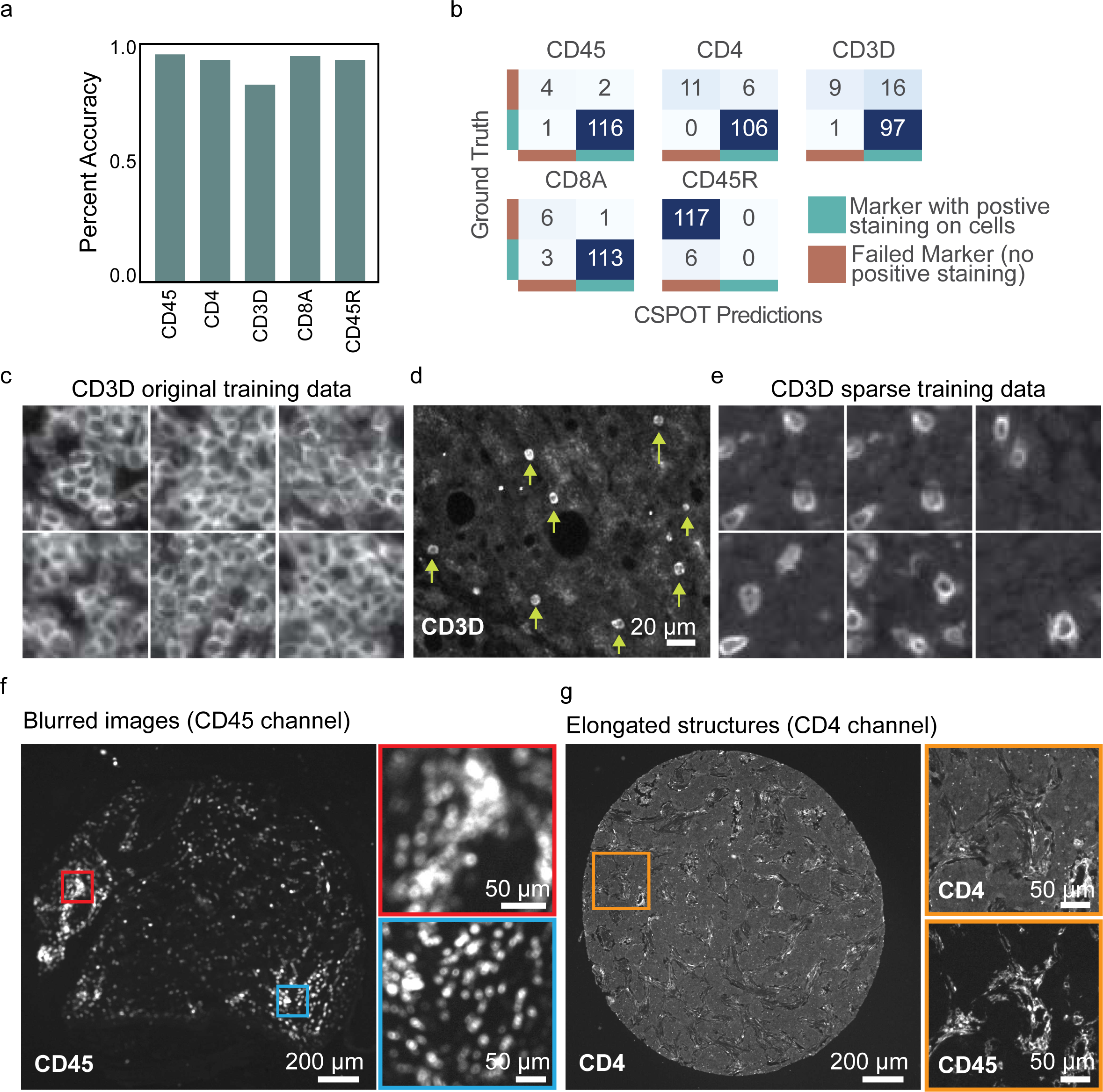
CSPOT identifies failed markers across 32 tissue types. a, Bar plot depicting the percentage accuracy (y-axis) of CSPOT’s classification predictions of positive or negative marker staining (x-axis) across 123 samples sourced from 32 tissue types. b, Confusion matrix comparing CSPOT predictions with the ground truth and further grouped by markers. c, Sample thumbnails utilized to train the CD3D model. d, Image highlighting the sparsely distributed CD3D^+^ cells (yellow arrows), which stands in stark contrast to the training data used in (d). e, Example thumbnails used to train a new CD3D CSPOT model that incorporates sparsely distributed CD3D^+^ cells. f, Image of an instance where the CD45 model was unsuccessful due to image blur. The panels on the right show magnified views of the red and blue boxes. g, Image of an instance where the CD4 model was unsuccessful due to unexpected dendritic-like structures. Panels to the right represent a magnified view of the orange box, highlighting CD45^+^ CD4^+^ cells.

A related problem was observed in the case of the CSPOT model for CD45: in this case, discrepancies relative to human annotators were concentrated in an area of the TMA in which tissue focusing was suboptimal (**Fig. 4f**). Moreover, CD4^+^ cells with dendritic morphology were missed (presumably myeloid cells; **Fig. 2g**); this issue could potentially be resolved, as the initial training dataset exclusively included CD4^+^ T cells with a compact, round shape. These data demonstrate that CSPOT models learn morphology, not just intensity. Effective training requires cells that exhibit the full range of possible morphologies and types of neighborhoods, both densely packed and isolated. Often, this can most easily be accomplished by using different tissue types during training.

### Cell type assignment using CSPOT

Cell type calling involves the identification of specific cell types based on known patterns of antigen expression, most commonly differentiation markers. It is usually performed using data in the spatial feature table and is therefore subject to the same difficulties in gating described above (**Fig. 5a**). To assign cell types using CSPOT using scaled marker intensity values, we developed a hierarchical assignment algorithm that uses logical qualifiers such as *’positive’*, *’negative’*, *’all positive/negative’*, and *’any positive/negative’,* etc. Hierarchical assignment started with the most broadly expressed markers (e.g., CD45 for immune cells and ECAD for epithelial cells), followed by subcategorization into CD3^+^ T cells, CD20^+^ B cells, etc. (**Fig. 5b, Extended Data 5a,b**). If a cell does not satisfy any of the pre-defined conditions, it is labelled as ‘unclassified’. When we compared cell types assigned by CSPOT and human annotators across the four TMA cores, F1 scores ranged from 0.78 to 0.92 and average accuracy from 0.92 to 0.99, depending on sample and cell type. A confusion matrix revealed two cell types with the greatest discrepancy (**Fig. 5c**). In one case, ECAD^+^ KI67^+^ epithelial cells, CSPOT was more effective than initial human curation (as judged by visual re-review) due to the presence of KI67 staining artifacts (**Fig. 5d**). In the second case, CSPOT identified CD8A^+^ cells that had been overlooked during manual gating (**Fig. 5e,f; Extended Data 5c**). Thus, CSPOT can be used to perform cell type assignment with super-human performance in some cases.

**Fig.5:**
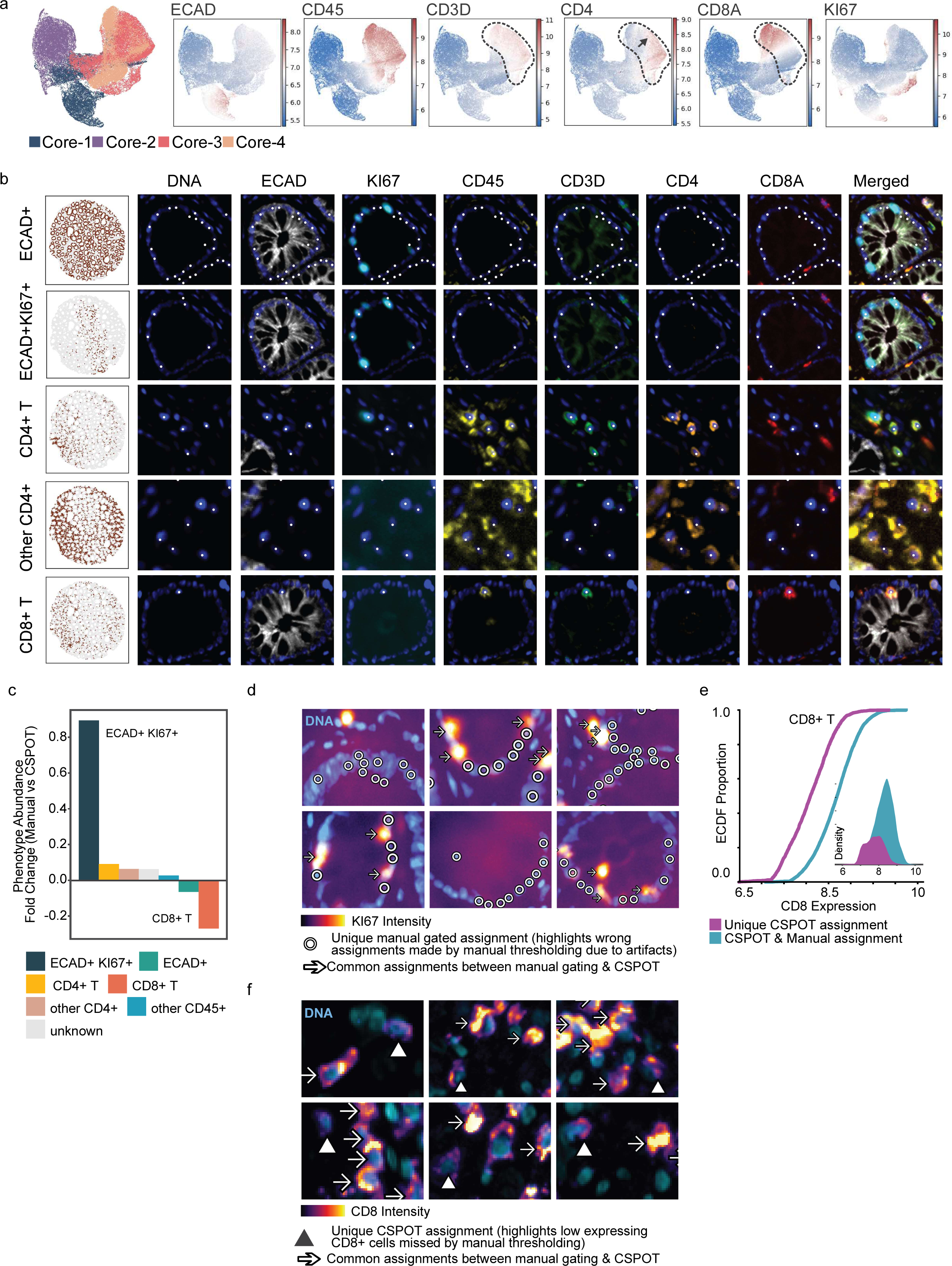
Cell phenotyping with CSPOT. a, The visualization of cores 1-4 using Uniform Manifold Approximation and Projection (UMAP). The cells are colored based on the sample (left image) and marker intensities (rest), with each dot representing a single cell. Dotted circles represent regions of CD3D expression likely corresponding to T cells. Arrow indicates a region of cells that are positive for both CD4 and CD8. b, Left-hand column shows an overall view of core-1 with cells mapped to their respective cell types (as predicted by CSPOT) based on spatial coordinates. Remaining columns display images from exemplary regions of core-1, highlighting the cell types identified by CSPOT. The dots on the images represent cells belonging to the cell type indicated on the y-axis. Middle columns display DNA (blue) and the single immunofluorescent marker indicated in top row; right-hand column displays merged image. c, Bar plot representing the fold change between the predicted cell types obtained by CSPOT and the ground truth cell types determined by manual gating. Each bar in the plot represents a different cell type, and the height of the bar represents the fold change between the predicted and ground truth proportions for that cell type. d, Fields of view of a sample with visible remnants of KI67 staining. Arrows represent cells that have been manually gated as KI67 positive and predicted to be positive by CSPOT (true positives). The hollow circles represent cells that have been manually gated as KI67 positive but are in fact false positives due to high background levels. DNA is shown in blue, and KI67 is represented as a heatmap relating to their intensity levels in the cells. e, An Empirical Cumulative Distribution Function (ECDF) plot that highlights the difference in CD8 intensity levels between cells that were missed by manual gating (purple) and those that were identified by manual gating (cyan). The inset plot shows the same data as a distribution plot, with CD8 intensity levels on the x-axis and density on the y-axis. f, Selected fields of view of a sample containing CD8^+^ cells. Arrows depict CD8^+^ cells with high intensity levels, which were identified by manual gating. Triangles indicate CD8^+^ cells with low intensity levels that were missed by manual gating and only detected using CSPOT. DNA is shown in blue, and CD8 is represented as a heatmap of the intensity level.

### A whole-slide test case: Identifying the proliferative structure of a colorectal cancer

To assess the performance of CSPOT on whole-slide images, we focused on identifying PCNA positive cells in a previously described 24-plex 3D CyCIF colonic adenocarcinoma (CRC) dataset^12^. PCNA is expressed in the early G1 and S phases of the cell cycle and serves as a marker of cell proliferation^24^. **Fig. 6a** presents two serial sections from this dataset, acquired under identical conditions, yet exhibiting significant variations in PCNA intensity levels. Owing to this staining variability, PCNA-positive and negative cells could not easily be distinguished using a single manually assigned intensity gate. When this was attempted, both false positives and true negatives were discernible (**Fig. 6b**) (e.g., when gates from one section - section 23 - were applied to another - section 12). In contrast, CSPOT effectively identified PCNA-positive cells (based on visual review) across all 25 tissue sections using a model trained from a single section (section 23), resulting in a substantial reduction in false positive cells (**Fig. 6c**). In this case, 9% of false positives were eliminated on section 12, and ∼17% of false negatives were recovered on section 22. Using CSPOT as a ground truth, we explored whether the manual gating led to significant false positive or negative PCNA cells in the CRC dataset. We found that when manual gates from one section (#23) were applied to the other 24 sections, 17 of the sections exhibited false positives and 2 exhibited false negatives, based on a 15% threshold (**Fig. 6d-f**).

**Fig.6:**
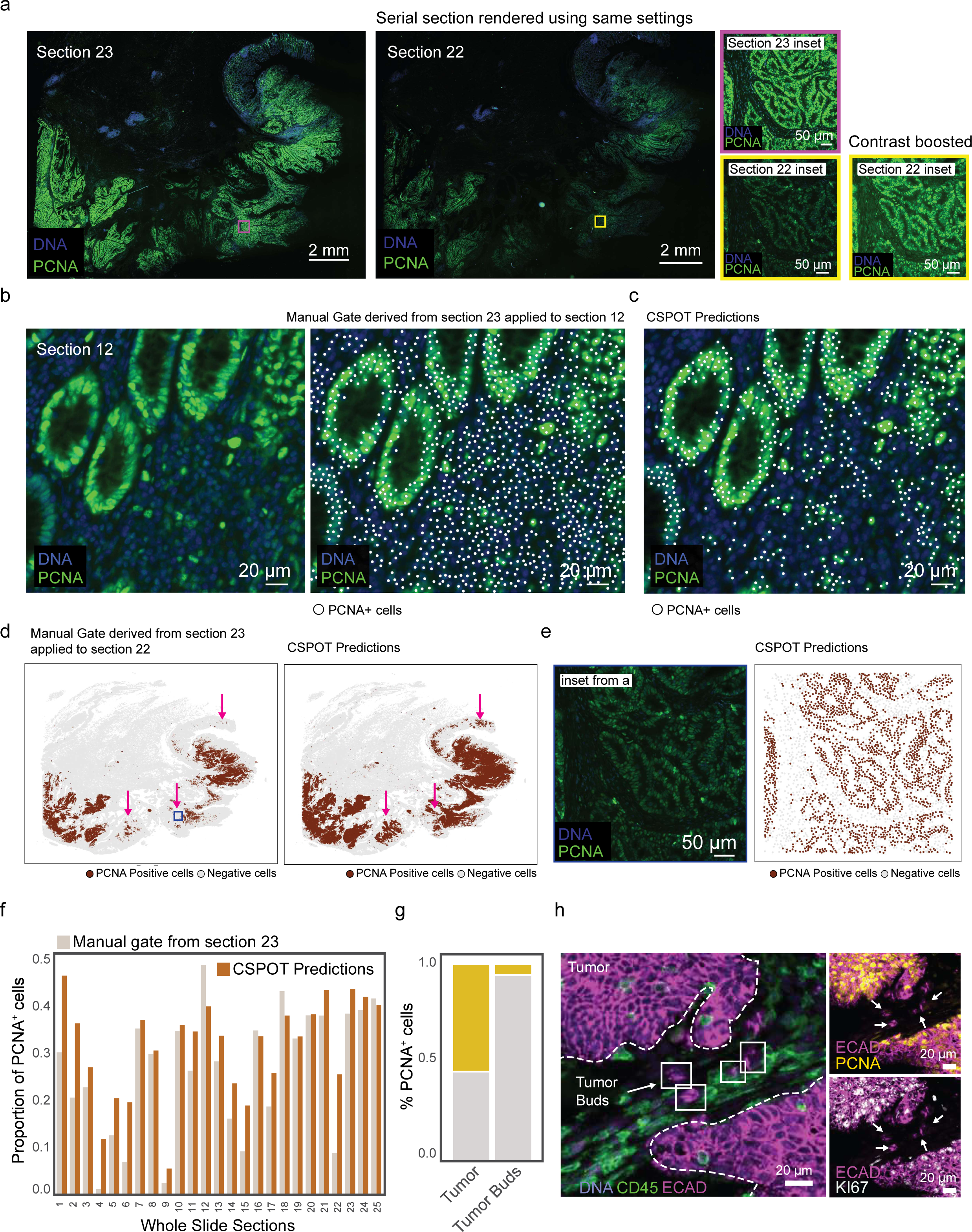
Application of CSPOT to whole slide images. a, Images of two adjacent sections of a colorectal tumor sample stained for PCNA, using identical rendering settings for both. The magenta and yellow boxes in the whole slide images are magnified on the right. Additionally, the inset of section 22 has been duplicated with a more contrasted rendering setting on the far right to make PCNA staining in that area more obvious. b, A field of view from section 12 shows PCNA expression (left), overlaid with PCNA^+^ cells identified through manual gating derived from section 23 (right). c, The same region as in b, but with PCNA^+^ cells as predicted by CSPOT. d, Scatter plot illustrating PCNA^+^ cells in section 22 identified by manual gating (left) and by CSPOT predictions (right); arrows indicate regions with significantly fewer PCNA^+^ cells, attributed to low intensity levels as shown in (a). e, An inset from section 22, shown in (a), displays low PCNA expression; the right plot indicates cells identified as PCNA^+^ by CSPOT. f, Bar plot showing the proportion of PCNA^+^ cells determined by both manual gating and CSPOT predictions; manual gating was conducted on one section and the same threshold applied to all sections. g, The proportion of PCNA^+^ cells determined by CSPOT in regions annotated as tumor and tumor buds across all 25 sections. h, A field of view highlights reduced PCNA expression (yellow) in tumor buds (indicated by arrows) compared to adjacent tumor regions (dotted areas). The subpanel to the bottom right indicate co-expression of KI67 (white).

After establishing the proliferative landscape using CPSOT, we examined PCNA expression in tumor cells and pre-annotated tumor buds, small clusters of tumor cells that are surrounded by stroma and associated with poor tumor outcomes and metastasis. These buds, annotated by a pathologist following the International Tumor Budding Consensus Conference (ITBCC) criteria, constituted approximately 0.002% (n=716) of the total cell population. A hallmark of these budding cells is their low proliferative index^25,26^. Thus, CSPOT was able to automatically and accurately assign proliferation status to a small population of diagnostically critical cells (**Fig. 6g**); hitherto this had only been possible using visual review (**Fig. 6h**).

## DISCUSSION

When applied to crowded cells in an immunofluorescence-based tissue image, intensity-based gating approaches effectively turn a microscope into a poorly performing flow cytometer: gating ignores the morphological information that is the most valuable aspect of a high-resolution image. This is also the information used by human observers to discriminate cells from one another and from background. Intensity-base gating becomes increasingly problematic as tissue images become more complex and densely packed and when multiple different tissue types must be analyzed in parallel. To overcome these problems, we have developed and tested a supervised ML-based framework for identifying cells in highly multiplexed tissue based on computation at both the pixel and single-cell level. CSPOT uses intensity and morphological information to automatically identify cells, despite variability in staining intensities, the presence of imaging artifacts (e.g., fluorescent smears and precipitates), and the confounding effects of high background. CSPOT also provides a morphology-aware approach to hierarchical cell-type calling that obviates the need for manual inspection of each image or marker and appears capable of identifying cells missed by human annotators. CSPOT is Dockerized and can therefore be incorporated into automated image processing pipelines such as MCMICRO^10^.

CSPOT has two ML components that work together, leveraging each other’s capabilities to enhance overall performance. An initial deep learning (DL) based module identifies true-positive and negative cells for a given marker, but with relatively poor recall of positive cells. A second module takes these cells as input and employs ML approaches such as a gradient boosting classification to more completely identify all positive and negative cells. Similar to other supervised methods, CSPOT performance depends on the accuracy and completeness of the training data. We found that effective training required ∼200 true positive and 200 true negative cells for each image channel. Models trained on one tissue type appeared to be effective on other tissue types, but it was important that representative cell environments and morphologies were included in training. For example, when we used training data from one tonsil tissue in which CD3D^+^ T cells were densely packed and round, we found that recall was poor with tissues in which CD3D^+^ cells were sparse; this problem could be overcome by additional training with data from images containing sparsely distributed cells. CSPOT therefore includes an optional neighborhood density parameter in the “*generateThumbnails*” function that can be used to generate training data arising from different types of cellular neighborhoods.

One limitation of the two-step ML approach in the current CSPOT implementation is that it can reintroduce anomalies that were eliminated by the DL algorithm. Ongoing work involves a one-step implementation eliminating the ML module. A second potential limitation of the current approach arises directly from our decision to analyze markers individually. This provides the greatest flexibility in the design of antibody panels but misses the opportunity to use complementary morphological data in other channels. Future work is being directed at assembling models from a set of near-universal markers (e.g., for basic immune cell types) or for antibodies with comparable staining patterns, for example those against common nuclear proteins or cell surface receptors.

## METHODS

### Generation of training data for model construction

CSPOT requires a one-time curation of training data for every marker of interest. This involves identifying true positive (POS) and true negative (NEG) cells for each marker of interest. Images containing the marker of interest (e.g., CD45) were segmented and quantified using the MCMICRO pipeline. From the quantified single-cell matrix, a two-mode Gaussian mixture model is fit to identify positive and negative cells. An optional parameter is to restrict the search within a user-defined intensity threshold (for example 2 to 12th percentile for negative cells and 88 to 98th percentile for positive cells, as used for this work). The centroids of these cells are used to cut out 64X64 pixel-sized thumbnails from the original image. The “*generateThumbnails*” function from the CSPOT package performs this function. A dedicated parameter “*restrictDensity*” has been implemented to allow users to discern positive cells that are sparsely distributed; it is to be used in scenarios where the markers do not exhibit any defined spatial pattern. Next, we sort through the thumbnails to remove any incorrectly assigned thumbnails. It is common for microscopy images to include artifacts, such as highly auto fluorescent cells, antibody aggregates, out-of-focus bright spots, illumination spikes, and dust particles, to be falsely detected as positive cells (purely based on high intensity). These need to be manually moved to the negative bin. Similarly, false negatives (true positive cells that have been incorrectly assigned to the negative bin) should be moved to the positive bin. As we are only interested in high-confidence examples, any ambiguous training examples are removed from both bins altogether.

#### CSPOT algorithm (Module 1: Deep Learning model construction)

The positive thumbnails are segmented using a combination of Gaussian blurring (with a standard deviation of 1) and Otsu thresholding^27^, to divide the image into foreground class (cells) and background class (rest). The entire image is considered as background for negative cells. We then randomly divide that data into training, validation, and test dataset with a 60:20:20 split ratio. Using the training and validation sets for each marker, a TensorFlow implementation of a 2-class UNet model^28^ is trained with 4 layers and 16 input features with a batch size of 16. We employed image augmentation (rotations and flipping), dropout, and L2 regularization to minimize overfitting. Importantly, we also employed on-the-fly intensity scaling augmentations to reduce the model’s reliance on just intensity values. For this purpose, thumbnail intensity values were either (1) raw and unaltered, (2) scaled to the entire image’s 99th percentile and minimum value, or (3) scaled to its own 99th percentile and minimum value. In each training epoch, thumbnails are randomly assigned to an intensity scaling class: 60% unmodified, 20% globally-augmented, and 20% locally-augmented thumbnails. All examples were loaded as 8-bit images. The training was performed with a learning rate of 0.0005 using an ADAM optimizer.

The created models are then applied to user-provided images in order to predict marker positivity. The probability maps for each pixel are returned as an image, which has the same dimensions as the original image. The CSPOT package’s "*generateCSScore*" function is used to compute and return the median probability score for each cell by using the pre-computed single-cell segmentation mask over the generated probability maps (referred to as the cspotScore).

#### CSPOT algorithm (Module 2: Predict all positive cells and negative cells)

The CSPOT algorithm’s second module utilizes the pre-computed marker intensity matrix and cspotScores as inputs. The intensity matrix goes through a pre-processing module where the marker’s intensity values are clipped to a range between 0.01 and 99.99th percentile to eliminate outliers, and then log-transformed. Our next step is to identify a set of bonafide positive and negative cells for each marker. To accomplish this, we fit a two-component Gaussian Mixture Model (GMM) on the cspotScore for each marker. The mean values of each of the distributions are then utilized to initialize GMM on the pre-processed intensity data. We found that utilizing the identified mean values to initiate the GMM greatly enhances the sensitivity to detect positive and negative cells, particularly in markers with skewed intensity patterns. Lastly, the cells that are recognized by both techniques are annotated as bonafide positive and negative cells.

Next, utilizing the bona fide cells identified in the preceding step, a Gradient Boosting Classifier is trained for each marker. The classifier’s input data includes the pre-processed intensity data for all markers present in the dataset for the bona fide cells, with the exception of certain markers such as background or DNA, as per the user’s discretion. The trained model is then employed to classify all positive and negative cells in the dataset for each marker. We then sort the data in ascending order based on the intensity level of each marker and find the point where positive cells first appear in the sorted data. This point is then used as a midpoint to re-scale the intensity data into a probability score between 0-1 by applying a sigmoid function. This method keeps the original distribution of the data while ensuring that cells with a probability score greater than 0.5 are always considered positive for the given marker and those with a score less than 0.5 are considered negative.

As a last step, the rescaled probability scores are passed into an anomaly detection module. This step aims to identify any highly-confident negative cells that have been misassigned as positive. Initially, we employ a Local Outlier Factor technique with 20 neighboring cells to identify negative cells for each marker. Afterward, we search for any positive cells that have been categorized as negative by this method. If any are found, we verify if they were classified as negative using the GMM-fitted cspotScores. If so, these cells are assigned to the negative class. In the end, the probability scores are adjusted to account for these changes. This is done by checking if any cells with a "negative cells" label appear after the defined midpoint (0.5). If any are found, they are given a value of 0.49. These typically tend to the imaging artifacts that have artificially high intensity values.

### Cell phenotyping algorithm

The rescaled probability scores are used for cell phenotyping. Briefly, the algorithm assigns phenotype labels to individual cells based on a sequential classification approach, following a tree structure. In the first step, cells are initially classified into large groups such as tumor (e.g., based on ECAD) and immune (CD45 intensity). In the second step, the immune cells are further divided into cell types such as T cells, B cells, etc. and in the following steps, these cell types are further divided into finer subtypes. The algorithm uses a user-defined relationship chart (phenotype workflow) between markers and cell types. Each cell is binned into a phenotype class based on the highest probability score of a given marker. If a cell does not express any of the markers (i.e., <0.5 for all markers), it is assigned to an unknown class. This approach is based on the assumption that the probability of a real signal, such as the intensity of a specific marker, would be higher than the signals that arise due to chromatic or segmentation artifacts, such as bleed-through of signals from one cell to another. The algorithm also uses logical operators such as "AND, OR, ANY, ALL" as parameters, in combination with "POS or NEG", to define the desired cell types. This allows us to fine-tune the classification and ensure that only cells that express the desired markers are classified into a specific phenotype class. If a cell type is defined by multiple markers, and one or more of those markers has a probability score less than 0.5, it is auto-labeled as "Likely-cell type." This is useful for detecting anomalies or, in some cases, tumor cells where they are defined by loss of unknown markers. Overall, this algorithm in combination with the scaled probability scores enables rapid and highly customizable classification of individual cells.

### Exemplar dataset for benchmarking CSPOT

A publicly available multi-tissue tissue microarray dataset^10^ was obtained to test CSPOT. We identified samples within the dataset that presented significant issues that often hinder the automated processing of multiplex imaging without human intervention. These obstacles include: (a) A marker that failed to stain cells in all images of the dataset, but still shows high levels of background fluorescence, (b) Images or markers that exhibit significant imaging artifacts such as antibody aggregates, unwashed antibodies, etc, (c) Cells with shapes that deviate from a circular geometry, (d) Antibodies that are able to stain some images, but not others, due to the absence of the cell type or other confounding factors, (e) Images where a majority of the cells are either positive or negative for a specific marker, resulting in skewed intensity distribution, (f) Examples of markers that stain the nuclei or cell surface. Four cores (core-1: Non-neoplastic colon, core-2: Mesothelioma, core-3: Seminoma, core-4: Mesothelioma) and 7 markers (ECAD, CD45, CD4, CD3D, CD8A, CD45R, KI67) were identified for single-cell assessment.

### Evaluation of published methods with artifactual data

We assessed the efficacy of two cell phenotyping methods, CELESTA^16^ and STELLAR^15^, using a normal kidney cortex core from the tissue microarray that showed unspecific binding of CD45R. The cell-type signature matrix for CELESTA was created to distinguish between two cell types: epithelial cells based on ECAD positivity, and immune cells based on CD45R positivity. The program was run using default settings as per the author’s instructions. STELLAR requires that a deep learning model must be trained on a dataset that has been previously phenotyped. In our study, we employed a Non-neoplastic colon core to identify epithelial and immune cells based on ECAD and CD45 intensity, which was used to train the STELLAR algorithm. Cells that did not fall into either category were excluded from the training process. Subsequently, the trained model was employed to predict epithelial and immune cells in the normal kidney cortex core. However, CD45R was used in place of CD45 during the prediction stage. The expected outcome was that there would be no immune cells as CD45R failed.

### Evaluation of other imaging modalities and whole slide imagine data

For the purpose of this demonstration, a publicly available CODEX image of a human tonsil was utilized^10^. The pre-computed segmentation mask and single-cell table available in the public repository were employed for CSPOT analysis. Among the markers used to stain the dataset, CD45 and CD8A matched the models that we had trained for; therefore, those models were applied to the image to predict the positivity of CD45 and CD8A. However, upon visual inspection, CD45 was determined to have failed as no CD45^+^ cells were visible. The COMET dataset utilized in this study was obtained in-house as part of a technology access program. The analyzed image is derived from a melanoma sample^29^, with two markers matching our previously trained models for CD45 and CD3D. We employed MCMICRO software to generate the segmentation mask and single-cell matrix, which was then utilized for CSPOT analysis. Finally, the whole slide imaging data used in this study was obtained from a publicly available colorectal tumor dataset^12^, where multiplexed imaging was conducted on 25 adjacent sections separated by an interval of 25 microns each. The pre-computed segmentation mask and spatial feature table available in the public repository were utilized for CSPOT analysis. See **Supplementary Table 1** for access information.

### Manual gating of the exemplar dataset

An OpenSeadragon-based visual gating tool (https://github.com/labsyspharm/cycif_viewer) was utilized to establish cutoff values that distinguish cells that express or do not express a specific marker. In short, cells that meet the user-specified threshold are dynamically highlighted in the image viewer, which is better than only using distribution plots for gating. Gating is done for each marker and image individually. Gating was done at three different time points, the average was taken and verified by a pathologist. These identified cutoff values were then used to rescale the single-cell data values between 0 and 1 so that values above 0.5 indicate cell intensities of the marker and vice versa using a sigmoid function. The adjusted single-cell data was used to classify cell types.

### Statistical analysis and figure creation

The Accuracy, Precision, and F1 scores were calculated using the *accuracy_score*, *precision_score,* and *f1_score* respectively within the *sklearn* package in python. Accuracy (Rand index) is defined as TP+TN / TP+TN+FP+FN; Precision is defined as TP/(TP + FP), and recall is defined as TP/(TP + FN). F1 score is defined as 2(precision × recall)/(precision + recall), where TP = True positive; FP = False positive; TN = True negative; FN = False negative.

Parts of **Fig. 1** were created using the Biorender (https://biorender.com). Multichannel overlay images were created using Napari.

### Generation of training data for model construction

CSPOT requires a one-time curation of training data. For this manuscript, we curated training examples for CD45, CD3D, CD4, CD8, ECAD (surface markers), and KI67 (nuclear marker) from a publicly available colorectal cancer dataset^12^ and a human tonsil image collected in house using whole slide Cyclic Immunofluorescence (CyCIF).

## DATA AVAILABILITY

See **Supplementary Table 1** for information about where to access these datasets and for the associated HTAN and Synapse identifiers. New data associated with this paper will be available through the HTAN Data Portal (https://data.humantumoratlas.org) and through Zenodo (doi.org/10.5281/zenodo.10055404).

## CODE AVAILABILITY

All codes related to CSPOT can be found at https://github.com/nirmallab/cspot and (doi.org/10.5281/zenodo.10055404). Detailed documentation and tutorials for running CSPOT are available at https://nirmallab.com/cspot/.

## ACKNOWLEDGEMENTS

This work was supported by the Ludwig Cancer Research and the Ludwig Center at Harvard (P.K.S., S.S.) and by NCI grants U2C-CA233280, R00CA256497 (A.J.N.), and U2C-CA233262 (P.K.S., S.S.). Development of computational methods and image processing software is supported by a Team Science Grant from the Gray Foundation (P.K.S., S.S.), the Gates Foundation grant INV-027106 (P.K.S.), the David Liposarcoma Research Initiative (P.K.S., S.S.), Emerson Collective (P.K.S.). S.S. is supported by the BWH President’s Scholars Award. We would like to thank Jia-Ren Lin, Zoltan Maliga and Juliann Tefft, and Jeremy Muhlich.

## AUTHOR CONTRIBUTIONS

AJN, CY and PKS developed the concept for the study. AJN and CY developed software and performed data analysis. SS provided specimens and pathology expertise. All authors wrote and edited the manuscript. SS and PKS provided supervision.

## COMPETING INTERESTS

PKS is a co-founder and member of the BOD of Glencoe Software, member of the BOD for Applied Biomath, and member of the SAB for RareCyte, NanoString, and Montai Health; he holds equity in Glencoe, Applied Biomath, and RareCyte. PKS consults for Merck and the Sorger lab has received research funding from Novartis and Merck in the past five years. The other authors declare no outside interests.

---

**Extended Data Fig. 1:**
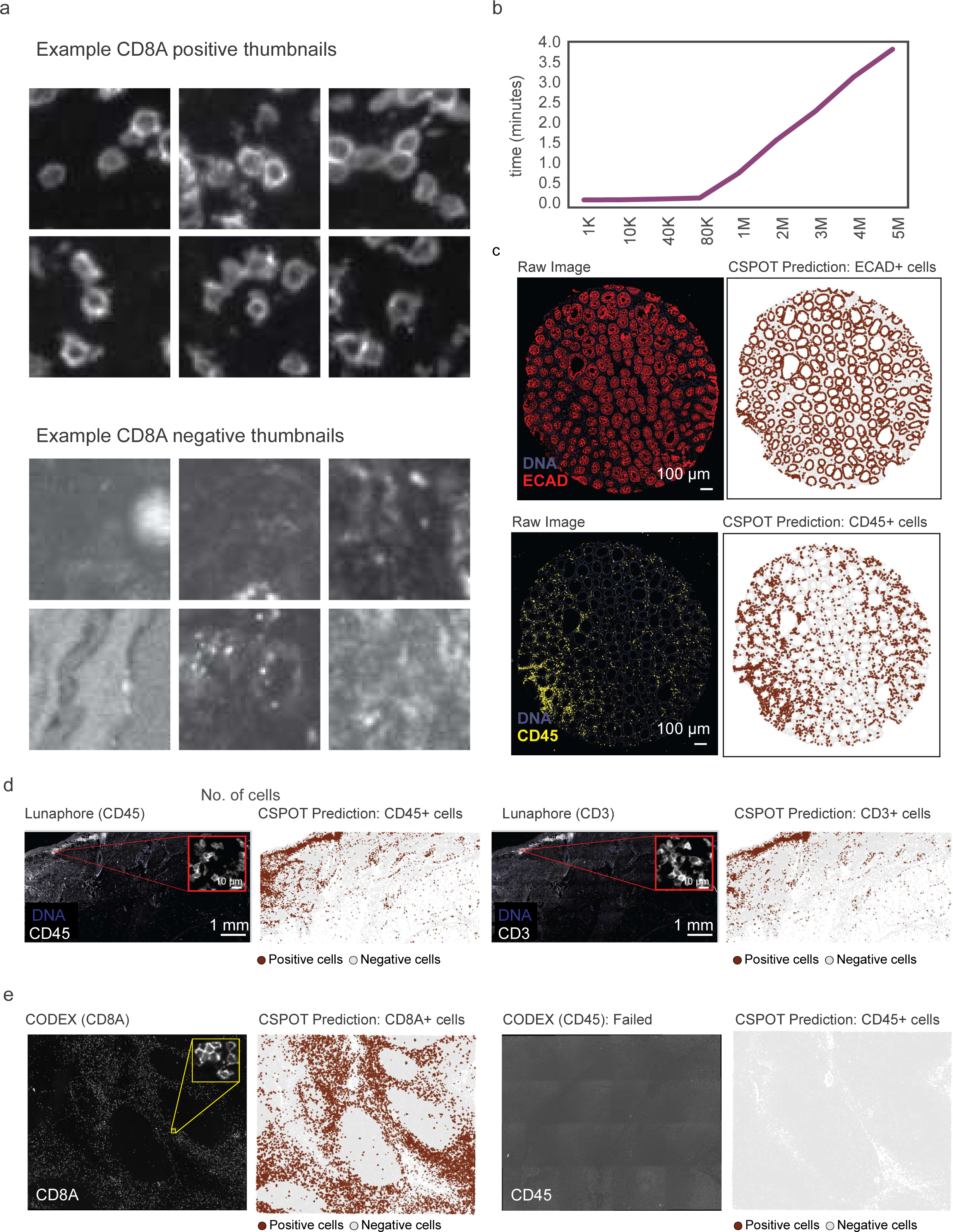
Testing CSPOT on other imaging modalities. a, Examples of training data that were used to train the CSPOT model for CD8A. Each thumbnail in this figure depicts a positive or negative example of the appearance of a CD8^+^ cell. b, Line plot showing the time (y-axis) required to run the CSPOT algorithm (single-cell level computation part of the algorithm) on varying image size/ numbers of cells (x-axis). The plot demonstrates the relationship between the number of cells and the computation time required by the CSPOT algorithm to perform prediction. c, The left image shows the ECAD and CD45 staining of a colon sample, while the right dot plot represents that were predicted to be positive for ECAD and CD45 respectively by CSPOT. d, The COMET image depicts the staining patterns of CD45 and CD3 proteins in the sample, along with the corresponding CSPOT predictions for these markers. Insets show a zoomed in view for a few cells. e, The CODEX image shows the staining patterns of CD8A and CD45 proteins in the sample, along with the corresponding CSPOT predictions for these markers. Inset shows a zoomed in view of a few cells.

**Extended Data Fig. 2:**
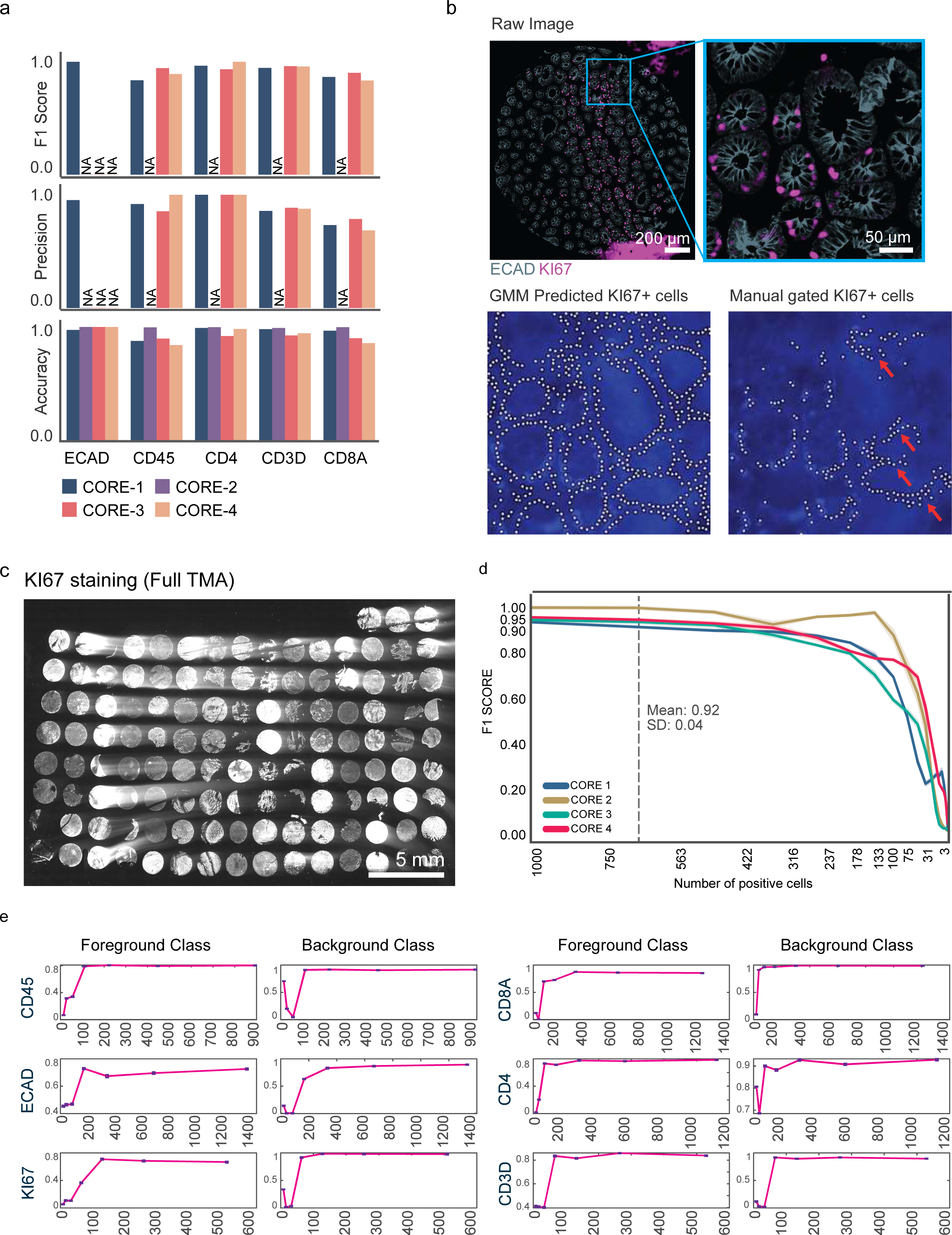
CSPOT accuracy and sensitivity. a, Bar plots for each marker in four samples (core1-4), representing the accuracy, precision, and F1 scores. b, The upper panels reveal residual KI67 staining with elevated background signal. The inset provides a closer view of a region containing actual KI67^+^ cells. The bottom panels represent cells that were identified as KI67^+^ either through the GMM method or manual gating, highlighting substantial number of false positives. c, Residual remnants of KI67 (white) staining across the entire tissue microarray. d, Line plot demonstrating the relationship between the number of positive cells present in the image (x-axis) and the resulting F1 score (y-axis). The graph indicates the minimum number of positive cells required to achieve a high F1 score. e, The figure displays line plots for seven markers, showing the number of thumbnails used for model training on the x-axis and the intersection-over-union (IoU) between prediction and ground truth on the y-axis. The plots demonstrate the relationship between the number of thumbnails used for model training and the accuracy of the model’s predictions, as measured by the IoU metric.

**Extended Data Fig. 3:**
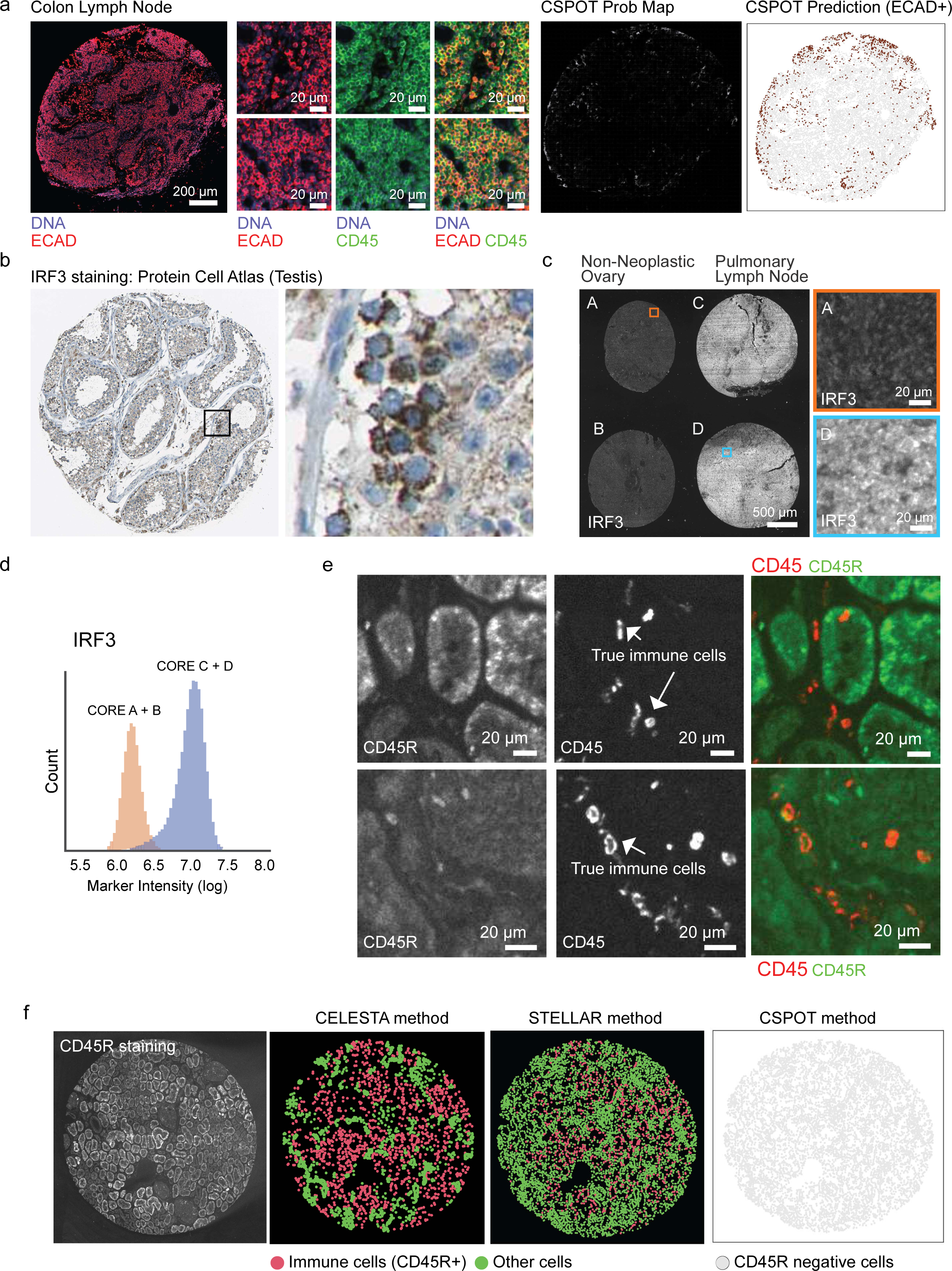
Uneven staining patterns impede automated image analysis. a, The image displays the staining of ECAD (red) and CD45 (green) in a sample taken from a colon lymph node. The smaller subplots provide a closer look at the samples. The aim is to draw attention to the ECAD’s non-specific binding on immune cells that are CD45^+^. The middle plots depict the probability maps generated by CSPOT at the pixel level and the right most plot depict the final CSPOT predictions for ECAD^+^ cells. b, The image, obtained from the Human Protein Atlas portal, illustrates the staining pattern of IRF3 in a testis sample using immunohistochemistry (IHC) staining. c, Tissue cores (non-neoplastic ovary and pulmonary lymph node) stained with IRF3 reveal differential background staining. The insets show magnified views of specific regions within the tissue sample. d, Intensity distribution plot of IRF3 from cores shown in (c). e, The top panel display a magnified view of a kidney sample, indicating the non-specific binding of CD45R. The structures stained with CD45R are considerably larger than actual immune cells stained with CD45 shown in the lower panels. f, The image on the left highlights a failed CD45R staining pattern. The middle plot shows cell phenotyping using the CELESTA method, which incorrectly assigns high CD45R background regions as immune cells. The plot on the right demonstrates the same analysis using the STELLAR method.

**Extended Data Fig. 4:**
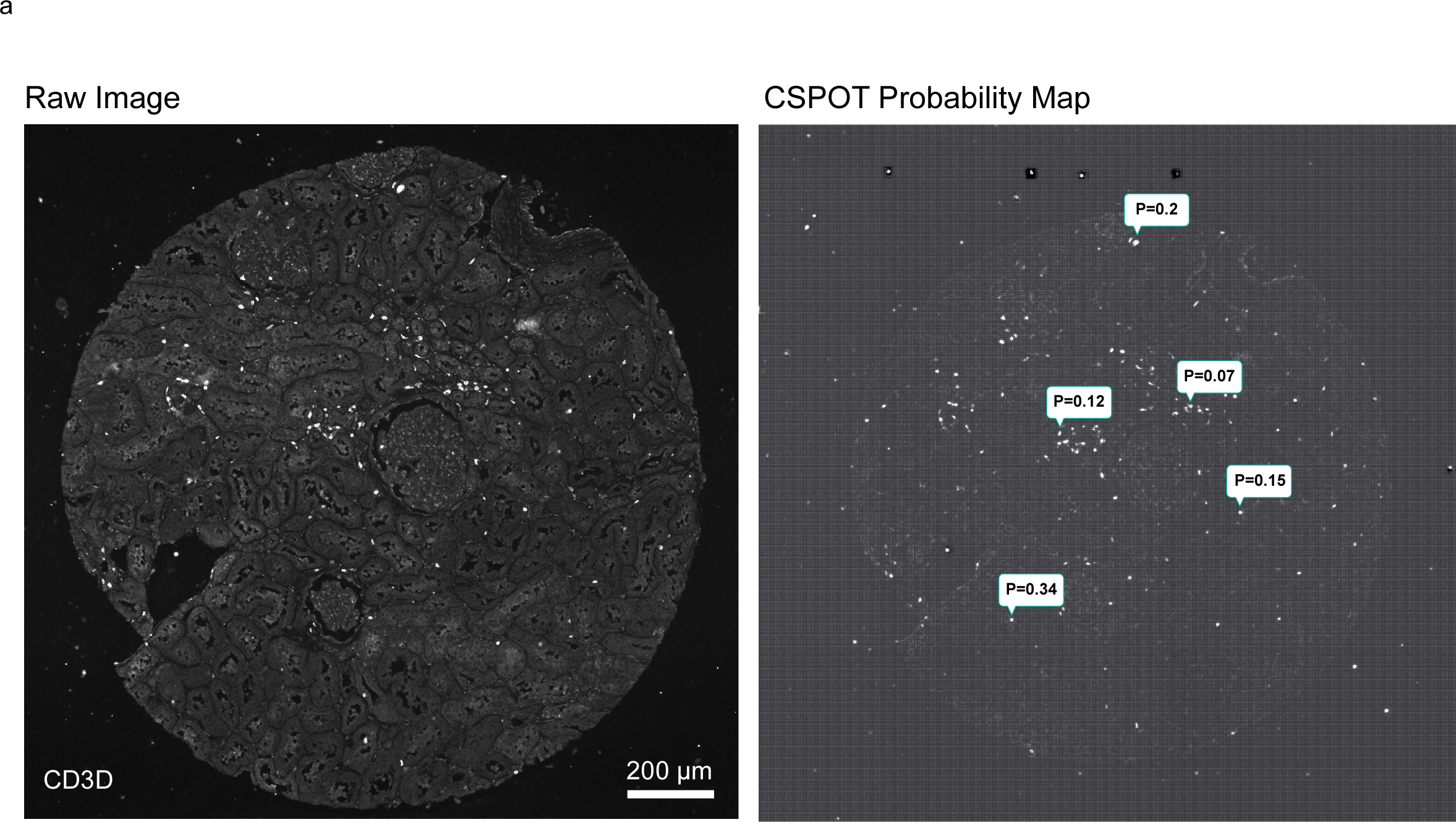
Failure of CSPOT. a, The left panel shows the raw image, where CSPOT failed to accurately classify the image as containing CD3D^+^ cells. The right panel shows the probability maps generated by CSPOT for the same image. The probability scores indicate that although CSPOT detected the presence of cells over the background, it displayed low confidence in detection.

**Extended Data Fig. 5:**
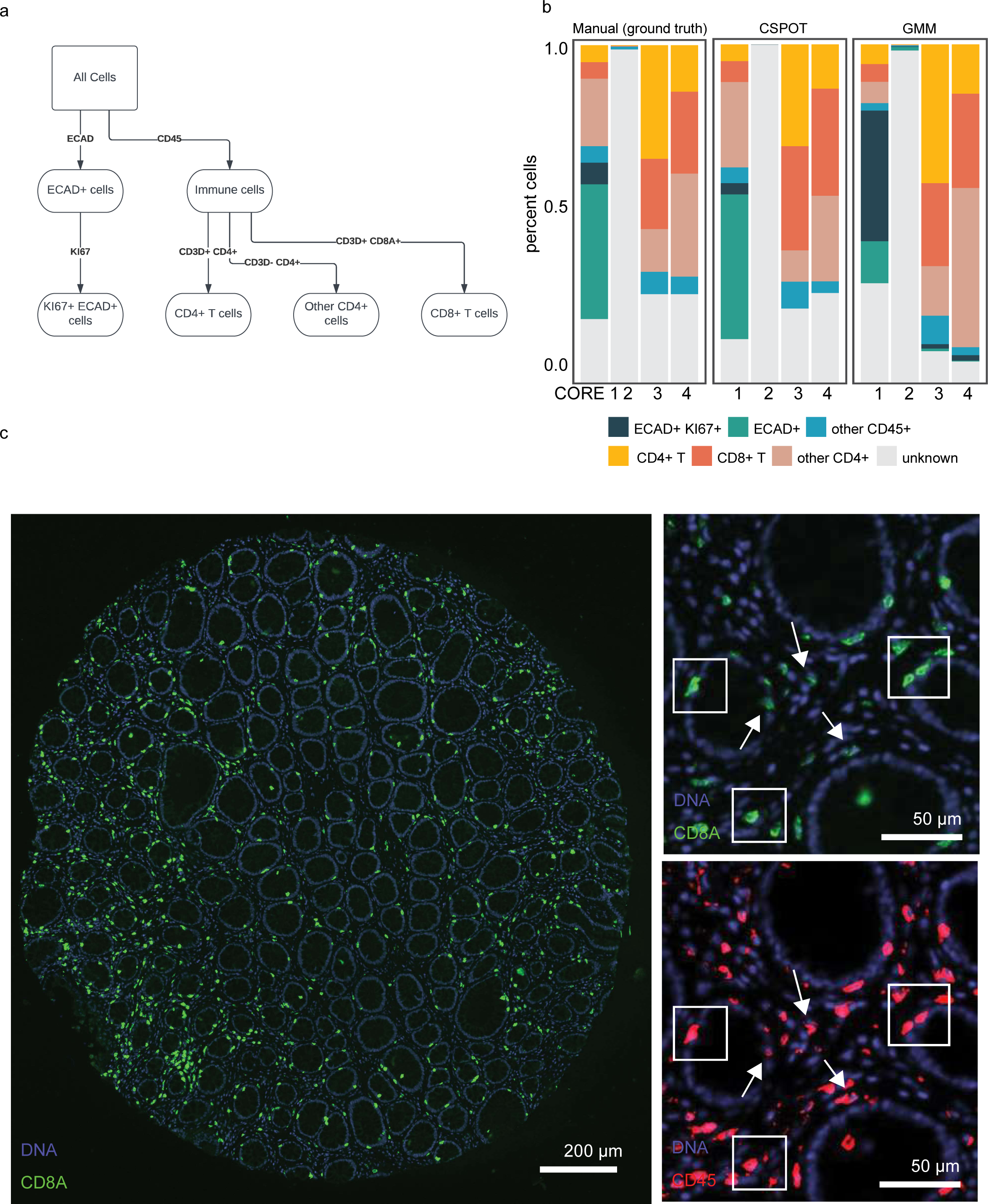
Cell Phenotyping with CSPOT. a, Flowchart used depicting the strategy used for cell phenotyping. b, Stacked barplot representing the proportions of cell types as determined by three different methods: manual gating, CSPOT, and GMM. Each bar in the plot represents the total proportion of cells for a given cell type, and the colors within the bar represent the proportion of cells classified by each method. c, Image highlighting CD8 intensity on a colon sample with a high signal (green) to background (black) ratio. The right plots display magnified views of the same image, indicating selected examples of cells with high CD8 (squares) and low CD8 expression (arrows). The bottom right panel displays co-expression of CD45 on the same cells.

