## Supplemental note for "Cell Spotter (CSPOT): A machine-learning approach to automated cell spotting and quantification of highly multiplexed tissue images"

Supplementary Note:

**Non-specific binding with variable intensity:** Alexa Fluor 647 conjugated antibodies against the IRF3 interferon regulated transcription factor failed to exhibit the expected pattern of cytoplasmic or nuclear staining in any core but varied sufficiently core-to-core. GMM on four adjacent cores (2 non-neoplastic ovary and 2 pulmonary lymph node samples) incorrectly classified the entire two non-neoplastic ovary samples (cores A and B) as negative and the two pulmonary lymph node samples (cores C and D) as positive for IRF3 (as shown in **Extended Data 3b-d**). CSPOT was not evaluated for IRF3 as no model had been constructed for this marker; however, this underscores how varying intensities of non-specific binding can confound automated approaches such as GMM. Similarly, Alexa fluor 647 -conjugated anti-CD45R antibodies are expected to stain B cells and subsets of T and NK but in kidney we observed staining to structures 5-10 times larger than typical immune cells reflective of non-specific antibody binding (as illustrated in **Fig. 3f,g, Extended Data 3e**) for a visual comparison with immune cells and evidence that this is not autofluorescence). The GMM algorithm, as well as other cell phenotyping methods like CELESTA and STELLAR, incorrectly classified these abnormally large, brightly stained structures as CD45R-positive cells (**Fig. 3h, Extended Data 3f**). In contrast, CSPOT was able to accurately identify these structures as negative for CD45R staining.
